# A multi-parent recombinant inbred line population of *Caenorhabditis elegans* enhances mapping resolution and identification of novel QTLs for complex life-history traits

**DOI:** 10.1101/443135

**Authors:** Basten L. Snoek, Rita J.M. Volkers, Harm Nijveen, Carola Petersen, Philipp Dirksen, Mark G. Sterken, Rania Nakad, Joost Riksen, Philip Rosenstiel, Jana J. Stastna, Bart P. Braeckman, Simon C. Harvey, Hinrich Schulenburg, Jan E. Kammenga

## Abstract

Local populations of the bacterivorous nematode *Caenorhabditis elegans* can be genetically almost as diverse as global populations. To investigate the effect of local genetic variation on heritable traits, we developed a new recombinant inbred line (RIL) population derived from four wild isolates. The wild isolates were collected from two closely located sites in France: Orsay and Santeuil. By crossing these four genetically diverse parental isolates a population of 200 RILs was constructed. RNA-seq was used to obtain sequence polymorphisms identifying almost 9000 SNPs variable between the four genotypes with an average spacing of 11 kb, possibly doubling the mapping resolution relative to currently available RIL panels. The SNPs were used to construct a genetic map to facilitate QTL analysis. Life history traits, such as lifespan, stress resistance, developmental speed and population growth were measured in different environments. For most traits substantial variation was found, and multiple QTLs could be detected, including novel QTLs not found in previous QTL analysis, for example for lifespan or pathogen responses. This shows that recombining genetic variation across *C. elegans* populations that are in geographical close proximity provides ample variation for QTL mapping. Taken together, we show that RNA-seq can be used for genotyping, that using more parents than the classical two parental genotypes to construct a RIL population facilitates the detection of QTLs and that the use of wild isolates permits analysis of local adaptation and life history trade-offs.

## Introduction

Determining how genotype-phenotype relationships are controlled is at the heart of genetics. Understanding how the relationships between traits, genotypes and environments are controlled is also crucial for traits relevant to the evolved context of the species [1, 2]. The identification and characterization of allelic variants associated with complex traits has been a major challenge in plant and animal breeding as well as disease genetics. Many complex traits vary in a continuous way across different genotypes of a species. It is this phenotypic variation that can be mapped to the genome using quantitative trait locus (QTL) analysis. Standard QTL mapping for many different species is based on recombinant inbred lines (RILs) derived from a cross between two genetically and phenotypically divergent parents. One of the many species that has extensively been used for exploring the genetics of complex traits is the bacterivorous nematode *Caenorhabditis elegans* [3, 4].

Genetic diversity between *C. elegans* populations on a local scale can be almost as diverse as on a global scale, with genetically distinct populations occurring within a few kilometers distance [5-9] and it is likely that both local adaptation and local competition between genotypes are critical for the species [1, 2, 10]. Most inbred mapping populations of *C. elegans* were derived from two globally distant locations, namely Bristol UK (N2 strain) and Hawaii (CB4856 strain) [4, 11-14]. These Bristol-Hawaii RIL populations have been very valuable for studying the genetic architecture of complex traits and the identification of genes underlying complex traits [15-25]. Even though other genotypes have been used in the construction of RIL populations, *e.g.* crosses between N2 and BO [26], N2 and DR1350 27], N2 and LSJ1 [28], JU605 and JU606 [29], MT2124 and CB4856 [30], JU1395 and MY10 [31], these additional RILs have only been used to address a specific question and thus their suitability to map QTLs for different types of traits is unclear compared to the work on the Bristol-Hawaii populations. Although other types of crossing strategies involving multiple lines 32 or panels of wild isolates have been reported, most of the work has been done on the Bristol-Hawaii derived RILs which only captures a subset of the phenotypic and genetic diversity present in *C. elegans.*

Inclusion of more than two parental sources of genetic variation and alleles captures more genetic variation and allows for more precise mapping and identification of potential regulatory variants of complex traits 33. An alternative to the conventional two-parental genetic mapping strategies is the development of Multiparent Advanced Generation Inter-Cross (MAGIC) lines. The first of such populations was developed for *Arabidopsis thaliana* consisting of 527 RILs developed from 19 different parental accessions 34. Since then many more MAGIC populations have been developed for a range of species 35. Recently, a *C. elegans* multi-parental RIL population originating from 16 wild-types 32 was characterized 36. This RIL panel comprised 507 strains originating from the 16 wild-types after experimental evolution for almost 200 generations and covered about 22% of single nucleotide polymorphisms (SNPs) known to segregate in natural populations [32, 36].

Here we report the construction and analysis of a multi (4) parental recombinant inbred line (mpRIL) population for *C. elegans*. The 200 mpRILs are derived from an advanced cross between four wild-types: JU1511 and JU1941 isolated from Orsay (France) and JU1926 and JU1931 isolated from Santeuil (France) (kindly provided by MA Félix, Paris, France; 8). The RILs were SNP genotyped based on RNA-seq data. We used the SNP genotyped lines for mapping QTLs for the following *C. elegans* phenotypes: length, width, length width ratio, volume, lifespan, lifespan during dietary restriction, heat-shock survival, oxidative stress, occurrence of males, the developmental speed on the food sources *Escherichia coli* OP50 and *Erwinia rhapontici* and population growth on *E. coli* OP50, *Erwinia rhapontici*, *Sphingomonas sp.*, non-pathogenic *Bacillus thuringiensis* strain DSM-350E, and the pathogenic *Bacillus thuringiensis* strain NRRL B-18247. We aimed to measure a range of traits under different bacterial food conditions and abiotic conditions that, to a certain extent, reflect natural conditions [8, 37, 38]. For all these traits heritable variation and QTLs were found. Here we present, a new multi-parental recombinant inbred line population and show the distribution of genetic variation, recombination, trait variation and identified quantitative trait loci and so show the effects of local genetic variation on phenotypic traits.

## Results

### Developing a *C. elegans* multi parental recombinant inbred line population

To allow the four parental genomes (JU1511, JU1941, JU1926 and JU1931)8] to recombine, we set up a crossing scheme in which two pairs of wild isolates were crossed and both the obtained F1 populations were reciprocally inter-crossed (Figure 1; Supplemental Table 1). To enable crossovers on Chromosome X and to generate extra crossovers, the heterozygous F2 obtained from these initial crosses were further inter-crossed. To create homozygous genotypes, single worms were selected from the F2 as well as from the F2 inter-cross for 6 generations of single worm inbreeding. From these 383 lines, a population of 200 different multi-parental recombinant inbred lines (mpRILs) was randomly picked for mRNA sequencing to obtain the genetic variation in coding sequence.

**Figure 1:**
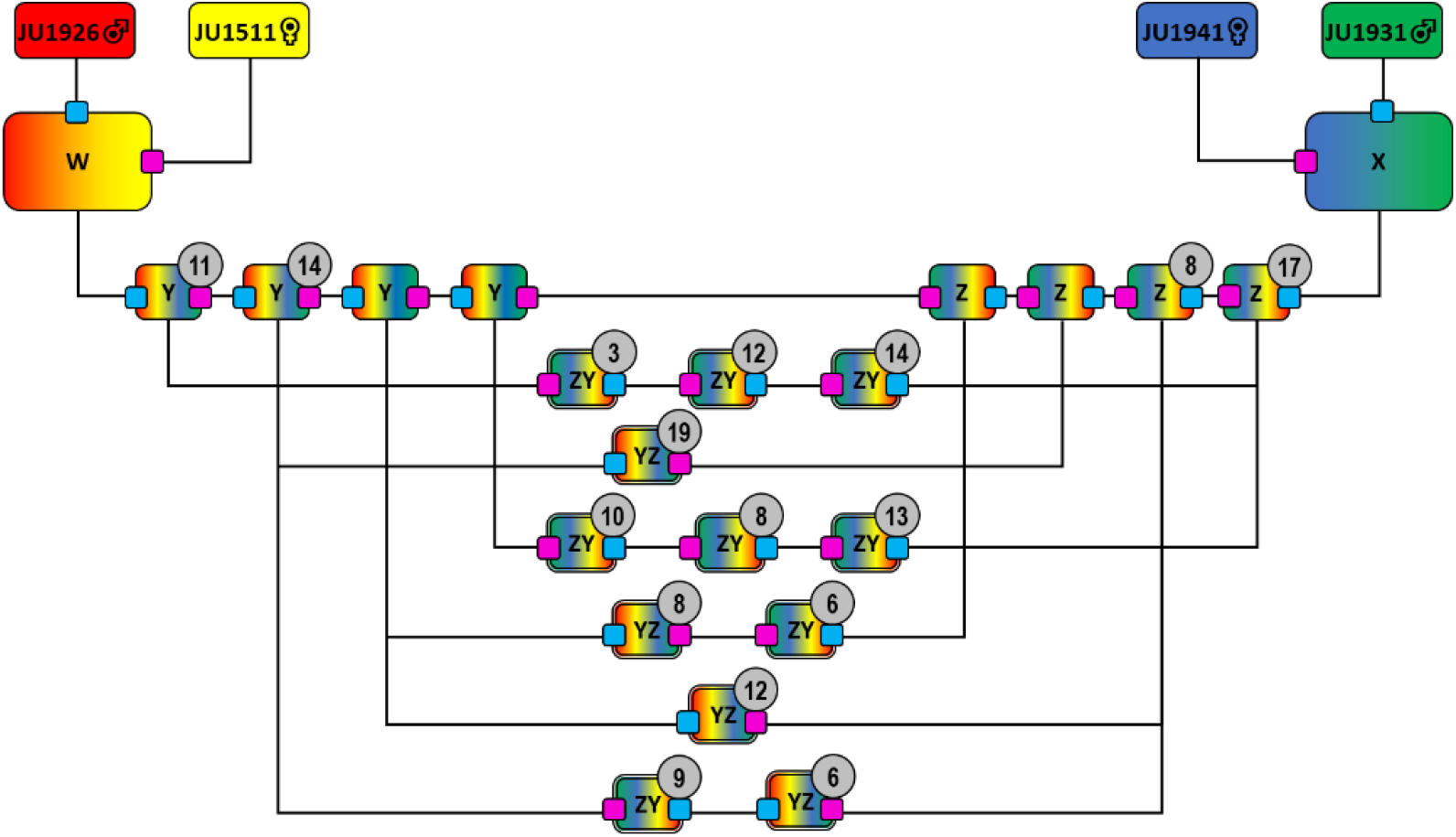
Crossing scheme used to make the four parental mpRIL population. JU1926 was crossed with JU1511 to create F1 population W. JU1941 was crossed with JU1931 to create F1 population X. Populations W and X were reciprocally crossed to obtain populations of genotypes with mixed genetic background from the four parental lines (Y and Z). Individuals from Y and Z where further intercrossed to obtain extra recombinations especially to break up the X-chromosome, which lacks recombination in the male. Numbers in circles show how many individuals were taken for single worm decent inbreeding for 6 generations to create mpRILs.

### Polymorphisms are not distributed equally

For genotyping of the 200 mpRILs we used the single nucleotide polymorphisms (SNPs) obtained from RNA-seq. We detected 8964 SNPs diverging in the coding sequences between the parental lines. The distribution of these SNPs over the genotypes can be grouped in 7 specific SNP distribution patterns (SDP): four patterns represent the four parental strains, three patterns are shared between two parental strains versus the other two parental strains. These are: SDP 12: difference between pair (JU1511/JU1926) and pair (JU1931/JU1941), SDP 13: difference between pair (JU1511/JU1931) and pair (JU1926/JU1941), and SDP 14: difference between pair (JU1511/JU1941) and pair (JU1926/JU1931). Importantly, SDP 14 therefore represents SNPs diverging between the two isolation sites and hence these polymorphisms may be informative of local adaptation. The SNP distribution differed between parents and the SNPs were unequally distributed across the genome (Table 1; Figure 2; Supplemental Table 2). For example, Chromosomes I, III, and V were more polymorphic in their coding sequences compared to Chromosomes II, IV, and X. Overall, Chromosome II was the least polymorphic and Chromosome III was the most polymorphic. Furthermore, we found regions where multiple SDP overlap (left arm of Chromosomes I, IV and V, right arm of Chromosomes I, IV, V and X and all of Chromosome III; Figure 2), which can potentially be used to reduce the number of candidate causal SNPs.

**Table 1:** SNP Distribution Patterns. Distribution of SNPs and alleles per chromosome I to X.

|  | Total | JU1511 | JU1926 | JU1931 | JU1941 | 12 | 13 | 14 |
| --- | --- | --- | --- | --- | --- | --- | --- | --- |
|  |  | Orsay | Santeuil | Santeuil | Orsay |  |  | O vs S |
| **Total** | **8964** | 2628 | 748 | 1651 | 2137 | 313 | 842 | 518 |
| **I** | 1916 | 288 | 10 | 75 | 1250 | 131 | 109 | 41 |
| **II** | 503 | 254 | 229 | 4 | 4 | 0 | 2 | 3 |
| **III** | 2231 | 979 | 16 | 217 | 476 | 72 | 281 | 158 |
| **IV** | 930 | 178 | 447 | 91 | 14 | 1 | 12 | 162 |
| **V** | 2152 | 104 | 11 | 1257 | 214 | 88 | 303 | 144 |
| **X** | 1232 | 825 | 35 | 7 | 179 | 21 | 135 | 10 |

**Figure 2:**
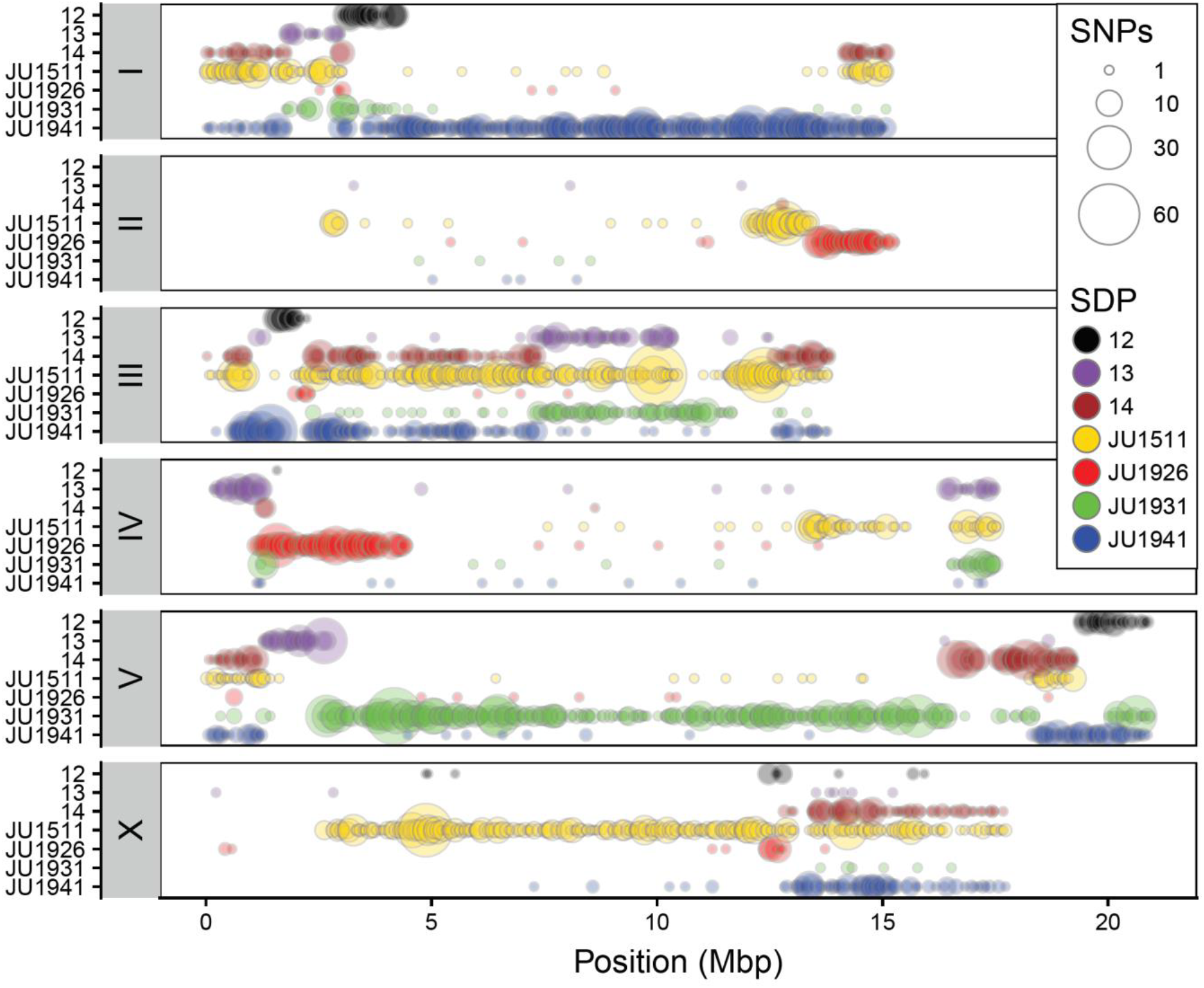
Genome-wide SNP distribution in the four parental genotypes. Circle size show no. of SNPs within 50K bin. Colours indicate SNP distribution patterns (SDP) as shown in the legend. These are: SDP 12: difference between pair (JU1511/JU1926) and pair (JU1931/JU1941), SDP 13: difference between pair (JU1511/JU1931) and pair (JU1926/JU1941), and SDP 14: difference between pair (JU1511/JU1941) and pair (JU1926/JU1931).

Most SNPs were strain-specific or diverging between strains from the two isolation sites. For example, unique SNPs on Chromosome I were mostly specific for JU1941 alleles, whilst those on Chromosome V were mostly specific for JU1931. For JU1511, unique SNPs were found on all chromosomes whereas the other parental genotypes have chromosomes almost completely lacking unique SNPs. Moreover, the genotypes from Orsay (JU1511 and JU1941) had more (>2000) strain-specific SNPs compared to those from Santeuil (JU1926 and JU1931). Almost 1700 SNPs were found in Orsay vs. Santeuil genotypes, whereas only 518 (~30%) were shared between genotypes from the same geographical location.

Across all chromosomes, both the left and right arms of the chromosomes (except for Chromosome III) were more polymorphic compared to the center of the chromosomes, a result matching that seen in previous work on *C. elegans* wild isolates [7, 11, 39]. Specific regions are very polymorphic between the four parental lines, such as the left arm of Chromosome I, all of Chromosome III, both arms of Chromosome V, and the right arm of Chromosome X. Long stretches of relative low SNP variation can also be observed, such as large parts of Chromosome II, the middle part of Chromosome IV, and left arm of Chromosome X. For the majority of the genome at least one parental genotype can be uniquely identified by individual SNPs. Overall, we conclude that SNPs in coding regions of *C. elegans* are unequally distributed over the genome and among genotypes, and that the chromosome arms are more polymorphic than the chromosome centers.

### Cross specific recombination in the mpRILs

The genetic map shows a highly variable frequency of recombination and introgression sizes (Figure 3A; Table 2; Supplemental Table 3 and 4). In total 1683 recombination events were found in the mpRIL population, with genome-wide allelic presence of the four parental lines (Figure 3B, Supplemental Figure 1). Up to four recombination events per mpRIL per chromosome were found, where one or two recombination events per chromosome was most common (Figure 3C). The average number of crossovers per mpRIL was 8.5 across all chromosomes and 1.5 per chromosome. Most recombination events were found on Chromosome III (348). As expected, due to lack of recombination of Chromosome X in males, the fewest recombination events where found on Chromosome X (233) (Figure 3D). Moreover, for Chromosome X almost 40% of the mpRILs showed no recombination. The recombination rate was on average once per 17 Mb, with a genome-wide mean introgression size of 5.0 Mbp (median: 3.1 Mbp). We observed a suppression of recombination across the centers of the chromosome with higher recombination rates at the chromosome arms (Figure 3E), consistent with previous work on *C. elegans* [7, 11, 40]. Considering the whole population, the genomic bins (loci) that can be individually investigated had a median size of 43 Kbp. (Table 2). The effective recombination rate useful for QTL mapping becomes larger as multiple SDP can be recombined by a single recombination event (Supplemental Figure 2). Including the SDP increased the effective recombination rate to approximately 57 per Mbp and 5686 in total. This does not affect the mapping resolution by making the QTLs substantially smaller yet it does affect which polymorphisms can be causal and therefore mapping in an SDP dependent manner affect the number of polymorphisms under investigation when looking for the causal gene or SNP.

**Table 2:**
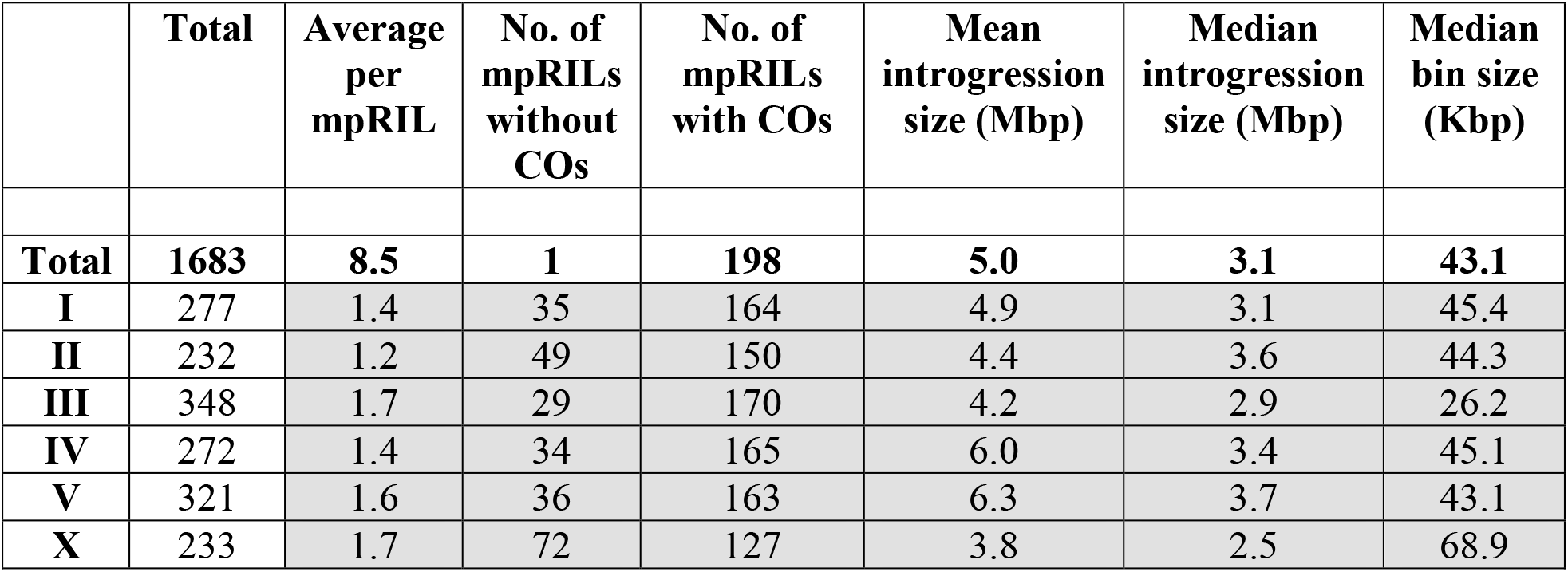
Crossovers per chromosome.

|  | **Total** | **Average**<br>**per**<br>**mpRIL** | **No. of**<br>**mpRILs**<br>**without**<br>**COs** | **No. of**<br>**mpRILs**<br>**with COs** | **Mean**<br>**introgression**<br>**size (Mbp)** | **Median**<br>**introgression**<br>**size (Mbp)** | **Median**<br>**bin size**<br>**(Kbp)** |
| --- | --- | --- | --- | --- | --- | --- | --- |
| **Total** | **1683** | **8.5** | **1** | **198** | **5.0** | **3.1** | **43.1** |
| **I** | 277 | 1.4 | 35 | 164 | 4.9 | 3.1 | 45.4 |
| **II** | 232 | 1.2 | 49 | 150 | 4.4 | 3.6 | 44.3 |
| **III** | 348 | 1.7 | 29 | 170 | 4.2 | 2.9 | 26.2 |
| **IV** | 272 | 1.4 | 34 | 165 | 6.0 | 3.4 | 45.1 |
| **V** | 321 | 1.6 | 36 | 163 | 6.3 | 3.7 | 43.1 |
| **X** | 233 | 1.7 | 72 | 127 | 3.8 | 2.5 | 68.9 |

**Figure 3:**
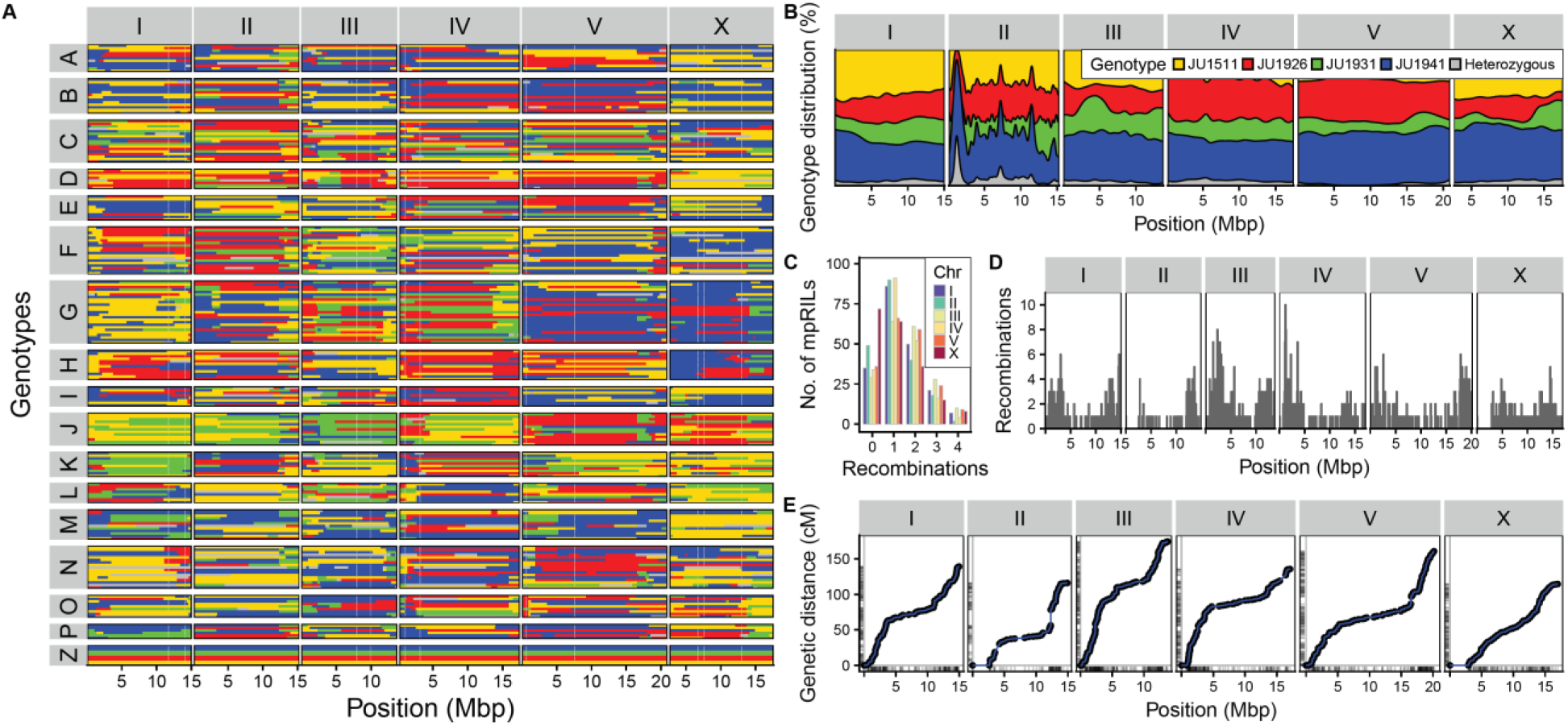
Parental background of the multi parental Recombinant Inbred Lines and recombination and allelic distribution per chromosome. **(A)** Colours indicate the parental background per genetic segment (x-axis) per RIL (y-axis) as estimated from the SNP Distribution Patterns. Chromosomes are in separate panels on the x-axis. mpRILs are grouped in according to their cross history. The parental lines are shown in group Z. **(B)** Recombination per chromosome. Chromosomes show on the x-axis. Number of recombination per RIL show on the y-axis. **(C)** Genome wide distribution of parental alleles. Colors indicate the percentage parental occurrence (y-axis) per genetic segment (x-axis) as estimated from the SNP Distribution Patterns. Chromosomes are in separate panels on the x-axis. **(D)** Recombination frequency per chromosome. **(E)** Genomic distance (x) vs genetic distance (y), rugs indicate the marker positions.

The allelic distribution was different between cross and inbreeding pools. The ratio of parental alleles shows a similar distribution across the chromosomes, except for Chromosome II (Figure 3B). Alleles from all four parents had a genome-wide representation, although JU1931 alleles occurred less frequently genome-wide and JU1926 alleles occurred relatively less frequently on Chromosomes I, III, and X. The allelic distribution was dependent on the specific cross and inbreeding pool (Figure1 and Supplemental Figure 1; Supplemental Table 1). In each specific pool the parental alleles display a cross-specific and chromosome-specific distribution, frequently showing absence of one or a few allele types (Supplemental Figure 1). Taken together, the whole population of mpRILs captures the genetic variation of the parental strains from which they were derived, perturbed by recombination.

### Phenotypic variation and heritability

As expected given previous work on *C. elegans* and the phenotypic variation between the parental lines 8, we observed substantial heritable phenotypic variation between mpRILs (Figure 4; Supplemental Table 5 and 6). Correlation analysis (Figure 5; Supplemental Table 7) across all phenotypic traits showed that timing of first eggs laid were highly correlated across different food conditions. This was also found for population growth, except growth on *Sphingomonas*. Body size and developmental phenotypes were also highly correlated. This shows that these phenotypes are likely to share a similar genetic architecture.

**Figure 4:**
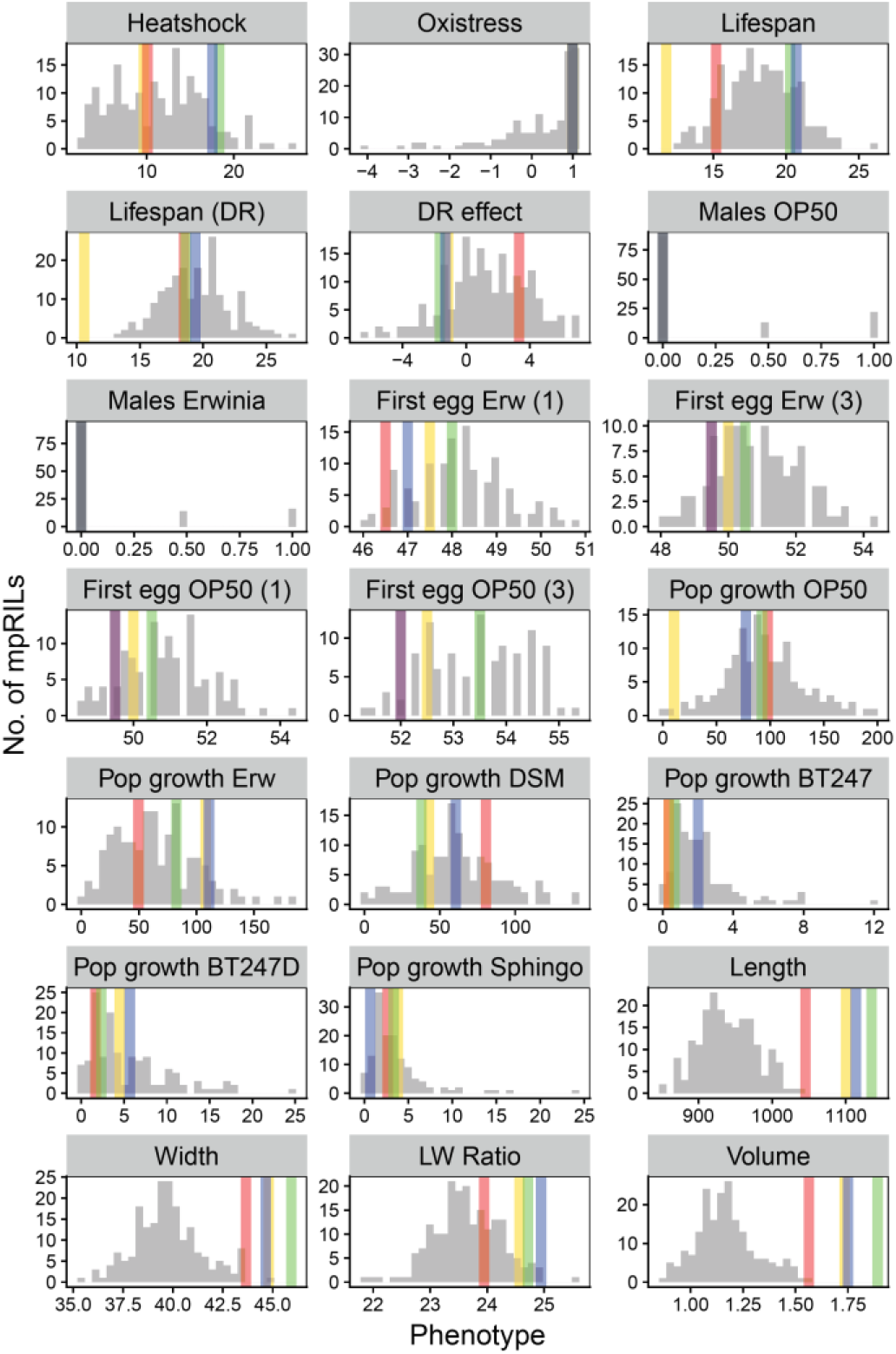
Phenotypic variation in the mpRILs. Distribution in the mpRILs are shown in grey, parental strains are shown in colour, yellow = JU1511, red = JU1926, green = JU1931 and blue = JU1941. Heat shock is average no. of animals dead per 50. Oxidative stress indicates activity. Lifespan is average lifespan on NGM in days. Lifespan (DR) is average lifespan on DR medium in days. DR effect is the difference in average lifespan between NGM and DR medium in days. Males OP50 and males Erwinia is occurrence of males on plates (0 = none, 0.5 = 1 plate, 1 = 2 plates). First egg Erw (1) is time in hours till first egg (1-10) for populations grown on Erwinia. First egg Erw (3) is time in hours till first egg (>100) for populations grown on Erwinia. First egg OP50 (1) is time in hours till first egg (1-10) for populations grown on OP50. First egg OP50 (3) is time in hours till first egg (>100) for populations grown on OP50. Pop growth shows worms per 5ul of culture. Length and Width in nm and Volume in nL, the parental values for these traits are taken from Volkers *et al.* 2013

**Figure 5:**
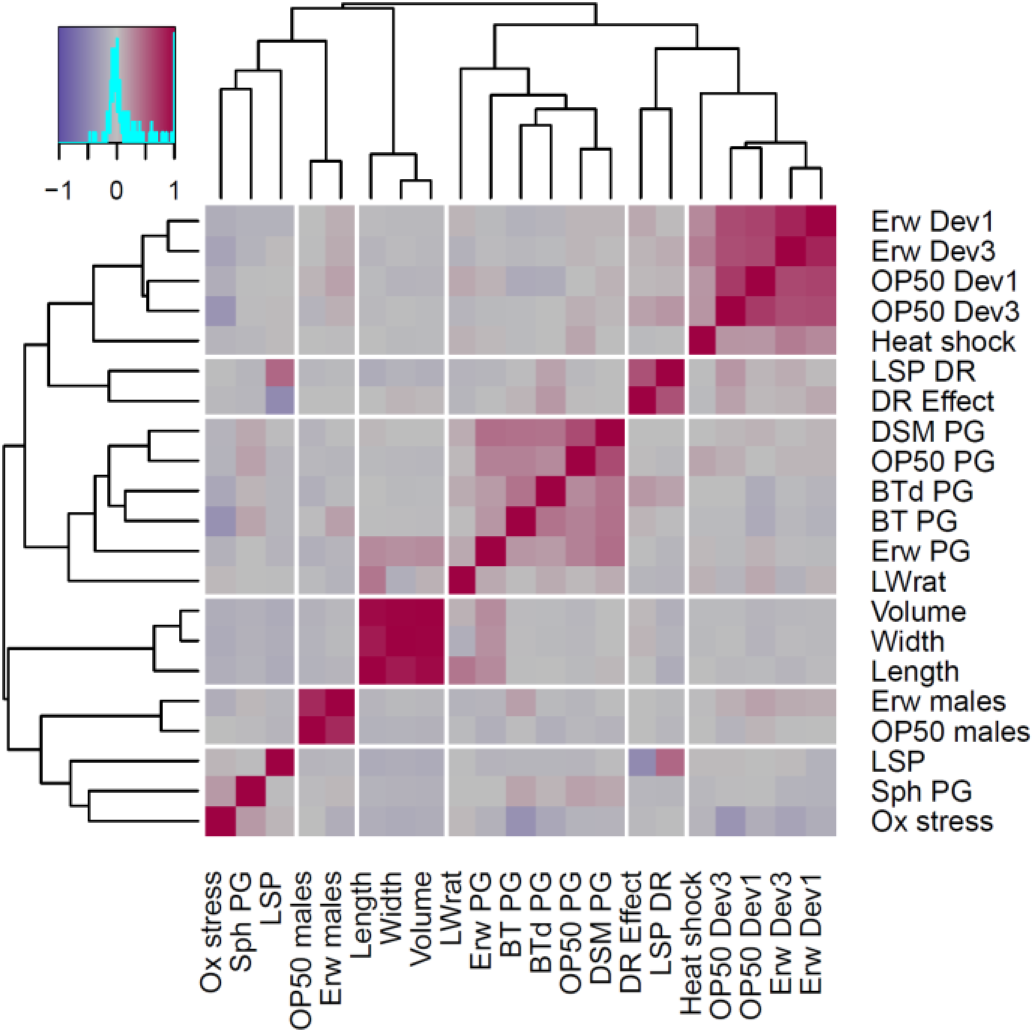
Correlations between traits. Pearson correlation in the mpRILs between traits.

Figure 4 shows the distribution of all measured phenotypes in the mpRILs and the parental wild-types. Average lifespan was 18 days (range: 13 to 26 d) and ~1 day longer under dietary restriction (DR) at 19 days (range 13 to 27 d). The overall effect of DR on lifespan was positive, but negative effects were observed for individual genotypes (~-6 days to ~+7 days) as previously found in *C. elegans* 41 and in mice 42. Heat shock (10 h at 35 °C) had a severe effect on the survival, on average ~11% (3 to 27) of the population survived 2 days after. Oxidative stress did not affect the average behavioral activity but did have an effect on individual genotypes (−4 to -1). Worms fed on *Erwinia* started laying their eggs earlier compared to worms fed on OP50. As previously found [8, 43] the average time of first egg laid on *Erwinia* was shorter (mean: ~51 hours; range: ~48 to ~54 hours) than on OP50 (mean: ~53 hours; range: ~51 to ~55 hours). The occurrence of males was similar on both OP50 and *Erwinia*, for most mpRILs no males were found, yet the genotypes that had males in the population did so on both OP50 and *Erwinia*. Population growth of the mpRILs differed strongly between worms fed different bacteria as previously found between wild-isolates 8. On average population growth was highest on OP50 (mean: ~93/5 μl; range: 0 to 197/5 μl), on *Erwinia* (mean: ~65/5 μl; range: 0 to 183/5 μl) and DSM (mean: ~61/5 μl; range 0 to 140/5 μl). Slow growth was observed on *Sphingomonas* (mean: ~3/5 μl; range: 0 to 24/5 μl) and on BT247 (mean: ~2/5 μl; range 0 to 12/5 μl). The mpRILs are also variable in length (mean: 945 μm; range: 848 to 1135 μm), width (mean: 40 μm; range: 35 to 46 μm), length width ratio (mean: 24; range: 22 to 26) and volume (mean: 1.2 nl; range: 0.9 to 1.9 nl).

The highest heritability of 83% was found for population growth on *Erwinia*, whereas the lowest of 55% was found on the most toxic concentration of BT247. Developmental speed for both OP50 and *Erwinia* showed high heritability of ~80% with many mpRILs having phenotypes beyond the parental phenotypic values (Supplemental Table 6). Most mpRILs had a slower development compared to the parents. Body size also showed high heritability (~80%), with length width ratio of ~70%. Population growth on the different bacteria showed variation in heritability, possibly linked to average growth rate on the specific bacteria. Transgression shows mpRILs beyond the parental phenotypes on both sides, yet for the growth on the BT247 strain transgressive mpRILs mostly produce better growth than the parents. Volume, length, and width phenotypes were all much higher for the four parents compared to the mpRILs.

Together the results show that ample phenotypic variation of complex traits can be found between the mpRILs and that these phenotypic differences are heritable. Genetic variation across the mpRILS causal for these different functional differences is likely to have major fitness effects.

### Quantitative trait loci

By applying quantitative trait locus (QTL) mapping using a forward co-factor selection approach we identified the loci associated with variation in the measured phenotypic traits (Figures 6 and 7; Supplemental Figure 3, Supplemental Table 8). The average QTL interval was 1.2 Mbp, median QTL interval was 0.88 Mbp, minimum QTL interval was 2.06 Kbp and maximum QTL interval was 7.7 Mbp. Most QTLs were found for the lifespan/stress traits (3-7 per trait) and together these explained between 32 up to 41% of the total trait variation observed. Of the 21 lifespan/stress QTLs most showed an allelic difference between JU1941 (7) or JU1511 (6) with the other three parental genotypes. The most significant QTL was found on Chromosome X at ~16 Mbp for the effect of oxidative stress which explained 20% of the variation. For developmental speed, 2 to 3 QTLs were found per trait and together explained 24% to 31% of the total variation per trait. Again, most of the 9 QTLs found for all developmental speed traits showed and allelic difference between JU1941 (3) or JU1511 (5) and the rest. Of all these QTLs a QTL found on Chromosome III around 12.3 Mbp for which the JU1511 allele shortens the developmental speed on *Erwinia* by almost 1 hour explained most variation (22%). Population growth traits showed between 1 and 3 QTLs, where the 3 QTLs for both *Sphingomonas* and DSM explained ~30% of the variation. The growth on other bacteria had less than 16% of the variation explained by the identified QTLs. The most significant QTL was found on chromosome V at 5.3 Mbp for the population growth on *Sphingomonas*, where the JU1931 allele increased population growth. For the body size traits 1 to 2 QTLs were found explaining up to 16% of the phenotypic variation, with the exception of the length width ratio for which 5 QTLs were found explaining 38% of the variation. Overall, we found that QTLs were mostly determined by JU1511 (21 QTLs) and JU1941 (14 QTLs) specific SNPs and relatively few by other alleles. Comparison of QTL locations for the different traits identifies no clear evidence of trade-offs between traits and only limited evidence for genomic regions affecting multiple traits (Figure 7). This suggests that selection can optimize traits without the kinds of trade-offs and associations that are seen in induced, laboratory derived, mutations.

**Figure 6:**
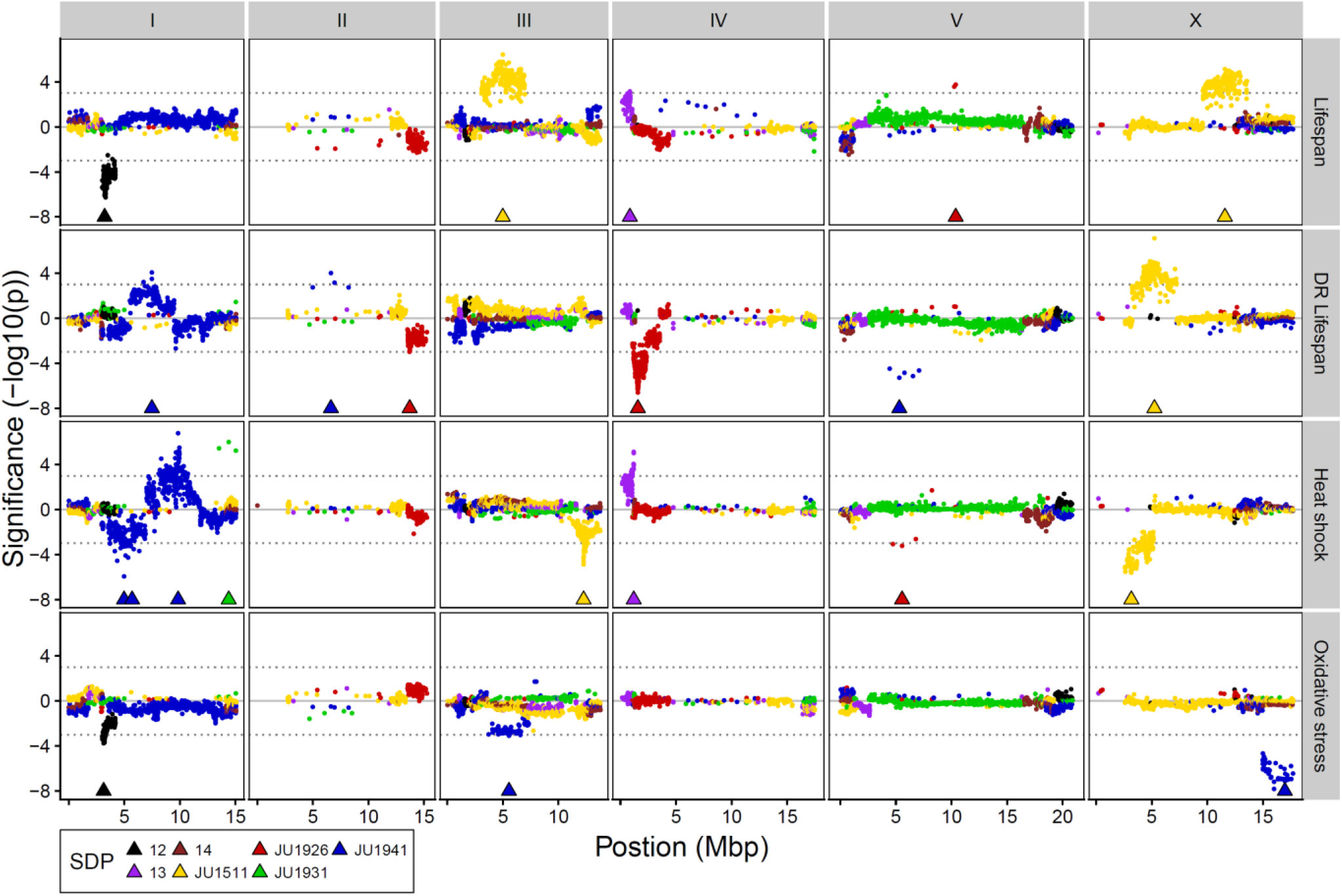
QTL profiles of lifespan and stress phenotypes. Genomic position on the x-axis against the significance on the y-axis. Triangles show the position of the cofactors used in final mapping model. Significance was multiplied by the sign of the allelic effect to show effect direction. Colours show SNP distribution patterns (SDP). These are: SDP 12: difference between pair (JU1511/JU1926) and pair (JU1931/JU1941), SDP 13: difference between pair (JU1511/JU1931) and pair (JU1926/JU1941), and SDP 14: difference between pair (JU1511/JU1941) and pair (JU1926/JU1931).

**Figure 7:**
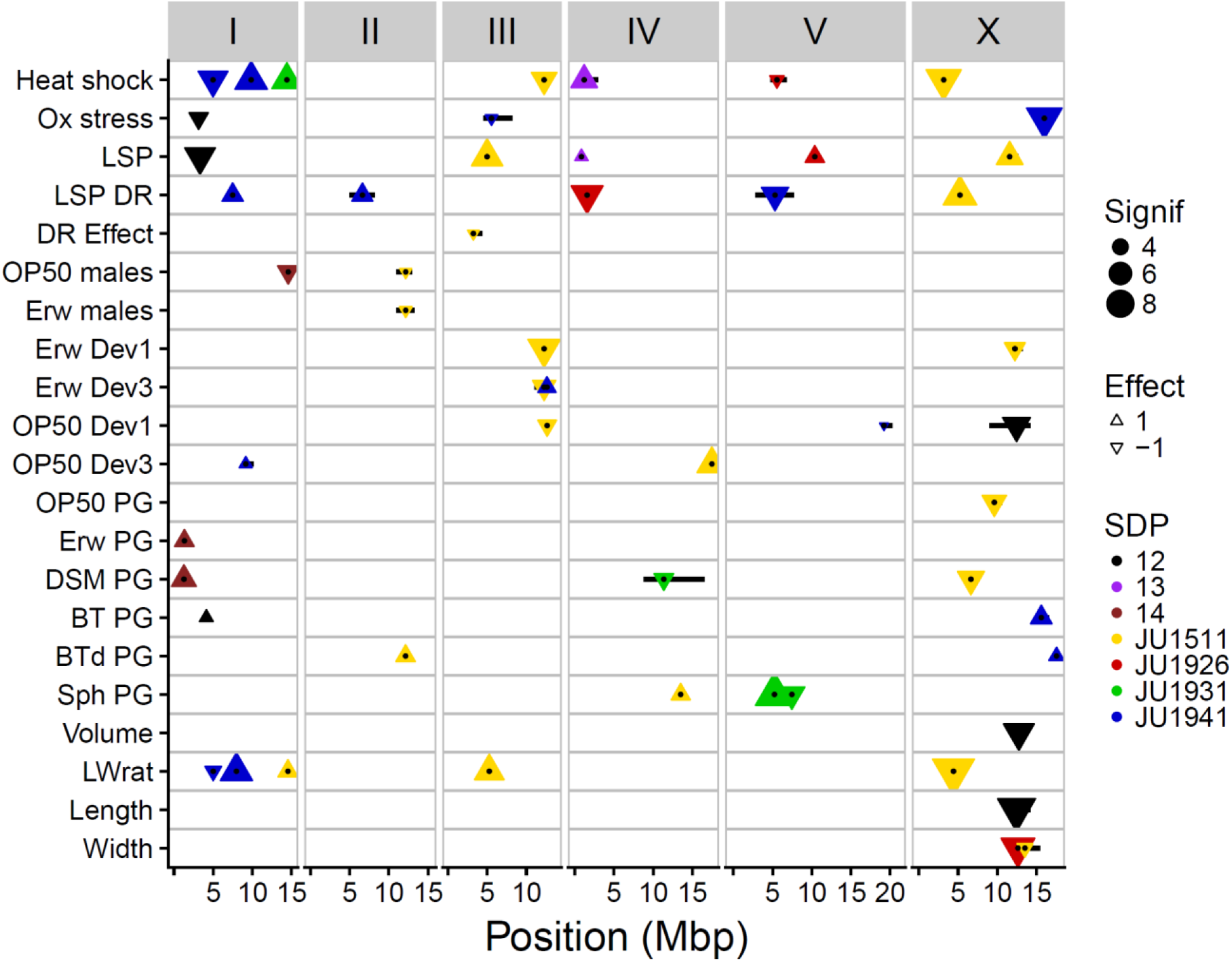
Genome-wide overview of QTLs. QTLs are show by the triangles, triangles pointing up show a positive allelic effect and pointing downwards show a negative allelic effect. Size indicate significance in–log10(p), colours show SNP distribution patterns (SDP). These are: SDP 12: difference between pair (JU1511/JU1926) and pair (JU1931/JU1941), SDP 13: difference between pair (JU1511/JU1931) and pair (JU1926/JU1941), and SDP 14: difference between pair (JU1511/JU1941) and pair (JU1926/JU1931). Black dots, show the exact location of the peaks and the black horizontal bars the 2-log10(p) drop QTL intervals.

## Discussion

We have developed a new *C. elegans* multi-parental recombinant inbred line population (mpRILs) derived from four wild-type strains capturing local genetic variation. This population of mpRILs complements existing RIL panels derived from two founders [11, 12, 26, 27, 29] and a multi-parent population derived from experimental evolution lines 36. The four founding parental wild-types originated from two different locations in France, Orsay (JU1511 and JU1941) and Santeuil (JU1926 and JU1931) 8. These strains were selected because of their genetic differences to each other. They also differed strongly compared to the two widely used strains Bristol N2 and Hawaii CB4856 8. These two canonical wild types can be considered genetic outliers because they differ extensively compared with many other wild type strains [4, 8].

Our mpRIL population is the first *C. elegans* population genotyped by SNPs in coding regions using RNA-seq. As RNA was isolated at a single time point (stage L4) during the life of the worm, these SNPs are limited to coding regions of genes that are expressed at that particular age. This limits the detection of SNPs to a temporally defined collection of expressed genes. There is a chance that this biases the detection of QTLs to SNPs of genes expressed at this point in the worm’s life. However, comparing the SNP distribution to genomic SNPs in *C. elegans*, we show that the SNPs based on RNA-seq are distributed according to expectation [7, 11]. Therefore, these SNPs are representative of the total genetic variation across the four strains. Hence, the QTLs detected are likely to be unbiased.

Recombination frequencies detected based on RNA-seq are also not likely to be biased when compared to those from two-parental RIL panels characterized by classical genetic markers or through genome sequencing. Yet, the average spacing between SNPs will be affected by RNA-seq genotyping compared to other populations genotyped by DNA sequencing. Our mpRIL population has a recombination frequency of 16.8 events per Mbp and a mean introgression size of

5.0 Mbp. When compared to the two-parental RIAIL population 11, the RIAIL population revealed a higher genome-wide recombination frequency of 36 per Mbp, most likely due to the advanced inter-cross design that was used to generate the RIAILs 11. Overall, we found 9000 SNPs with an average spacing of 11 kb compared to 1454 nuclear SNP markers with a spacing of 61,160 bp in the RIAIL population. A spacing of 1 SNP per 20 kb was reported for a multi-parental experimental evolution (CeMEE) panel, derived from 16 parental strains in *C. elegans* 36. As far as these results can be compared they show that all these populations are in the same range regarding SNP density, with an overall increase in SNP density for our mpRIL population.

Heritability values (50-80%) are relatively high for some traits compared to those previously reported (~ 20%) 44, yet similar to heritability values found for related fitness traits across other species. Moreover, the values reported in our study are in line with the ones reported by Noble et al. (2017) 36. It should be noted that heritability is a characteristic of a population and not of an individual trait. Therefore, heritability values are dependent on both the environmental and genetic circumstances of a population. Future studies with different RIL panels will help to understand to what extent heritability for specific traits can vary between populations of a species and whether and why there may be upper or lower boundaries for such measures. The range of inbred populations currently becoming available in *C. elegans* may be ideal to dissect variation in heritability in more detail.

We have investigated the genetic architecture of a range of complex traits and identified single and usually multiple QTLs underlying the observed variation. In total, QTLs with a JU1931 (4), JU1926 (4) or SDP 13 (2) allelic effect were found least frequent. QTLs with a JU1511 (21) and JU1941 (14) allelic effect were most abundant. This shows another advantage of using a mpRIL population over classical two parental RIL populations. Around 80% of the QTLs would have been found when just JU1511 and JU1941 had been used as parental strains for a RIL population. In contrast, only 30% of the QTLs would have been found using JU1926 and JU1931 as parental strains, thereby highlighting the advantages of a multi-parental RIL population. Furthermore, in the regions where multiple SDP overlap (left arm of Chromosome I, IV, and V, right arm of Chromosome I, IV, V and X and all of III) the number of potential causal SNPs can be reduced, as each QTL is mapped on a specific SDP. Estimated on coding SNPs this can be a reduction of 30-99% of the SNPs (Supplemental Table 8).

We compared the detected QTLs in the mpRILs to those reported from other studies, revealing identification of novel QTLs through the here developed mpRIL panel. For example, for variation in lifespan, in a two parent N2 x CB4856 IL population QTLs were detected on chromosomes II and IV 45 and in another study on chromosomes I, II, III, IV and X 14 while in the current study QTLs for the same trait were found on chromosomes I, III, IV, V, and X (Figure 7) – without any apparent overlap. For lifespan under DR, we identified QTLs on chromosomes I, II, IV, V; and X, again at different positions than in previous publications where QTLs for lifespan under DR were found in lines derived from N2 and CB4856 41. Population growth on the different bacteria, including the toxic Bt strains did not match with the QTLs found for leaving behavior on those same bacteria 46. QTLs for body size similarly differed between studies. Snoek et al (2014) detected a QTL for body size (length) on chromosome IV 45 whereas we found QTLs on other locations. For body size (volume) we found a QTL on Chromosome X whereas Gutteling et al (2007) reported QTLs on Chromosome IV and X 44. These examples emphasize that our new multi-parent RIL panel shows high power to identify previously undetected QTLs for complex traits.

Our findings support the idea that a substantial part of the variation in a range of complex traits in *C. elegans* is determined by a relatively limited number of QTLs of large effect rather than only by multiple QTLs of small effect. This is corroborated by other studies mapping complex traits in *C. elegans*. McGrath et al. (2009) identified 2 QTLs associated with digenic behaviour in response to environmental O_2_ and CO_2_ levels 20. Gaertner *et al*. (2012) mapped a set of few loci determining thermal preference and isothermal dispersion and found these loci to interact epistatically, explaining 50% of the total variation 47. Andersen et al. (2014) reported the detection of a few QTLs for lifetime fecundity and adult body size explaining up to 23% of the variation 15. In an extensive complex trait mapping study, Andersen et al. (2015) investigated the loci underlying fecundity and multiple body size traits 48. For fecundity, they found a single QTL on chromosome IV explaining 12% of the phenotypic variance. Comparing these results with the results obtained from the mpRILs, shows that multiple and different variation inducing alleles, even with a relatively small effect, are present within the total set of genetic variation in *C. elegans.* Moreover, the allelic effects might be dependent on the genetic background or epistatic interactions 36.

The advantage of mpRIL populations to the classical two parental RIL populations is that mpRILs capture a larger part of natural genetic variants in more combinations and hence covers these variants better. This is supported by similar multi-parent RIL studies in *A. thaliana* populations derived from 19 different parental accessions 34 and many more MAGIC populations were developed for different species, including mice 35. In our mpRIL population the polymorphisms show more patterns of segregations due to SNP distribution patterns between the parental strains, making candidate/causal gene selections more efficient. As each recombination event can break up 7 SNP distribution patterns, we found 57 informative breakpoints per Mbp, which can drastically reduce the number of candidate causal polymorphisms. These numbers suggest that our new panel may help to characterize the genetic architecture of complex traits at high resolution. Moreover, our developed four parent mpRIL population adds a new mapping tool for studying complex trait architecture in the model species *C. elegans* and complements existing RIL panels using 2 and 16 parents. In fact, it provides a straight forward alternative next to the more complex and less balanced CeMEE panel. Compared to the latter population, our mpRILs represent a relatively equal distribution of standing unperturbed local natural genetic variation as opposed to genetic variation partially derived from laboratory selection experiments 36.

Overall, multi-parent RIL populations have a higher number of informative SNP markers than the classic two parental RIL sets in a variety of organisms. We show that in our mpRIL population the number of QTLs is likely to be increased as well as the distinction of candidate causal SNPs and therefore resolution for genetic characterization of complex traits.

## Acknowledgements

We thank the compilers of WormBase for making it a versatile and important resource for *C. elegans*. We acknowledge financial support from the Deutsche Forschungsgemeinschaft to HS, grant number SCHU 1415/11 and project A1 within the CRC 1182. Furthermore, financial support from the NWO-ALW (project 855.01.151) to RJMV. The funders had no role in study design, data collection and analysis, decision to publish, or preparation of the manuscript.

## Author contributions

LBS, HS, SCH and JEK conceived the study. LBS, HN, MGS analyzed the data. RJMV, BPB, CP, PD, RN, JR, PR and JJS performed the experiments and/or provided phenotypic data. LBS and JEK wrote the paper with contributions of all authors.

## Methods

### C. elegans strains, culturing and crossing

*C. elegans* strains were cultured at 20 °C on OP50, unless specified otherwise for a specific screen or cross. For the construction of the multi-parental RIL population lines were crossed as described in Supplemental Table 1 followed by 6 generations of inbreeding. Males were induced by heat stress (4-6 h at 30 °C).

## RNA-sequencing

### RNA isolation

For each mpRIL and parental strain, worms were grown on two 6 cm dishes at 16 °C on OP50 and bleached at adult stage. Eggs were distributed over two 6 cm dishes and grown at 24 °C for 48 h after which the animals were rinsed of the plates and flash frozen in liquid nitrogen. We isolated RNA from these samples using the Maxwell^®^ 16 AS2000 instrument with a Maxwell^®^ 16 LEV simplyRNA Tissue Kit (both Promega Corporation, Madison, WI, USA). For isolation, the protocol was followed with a modified lysis step. In the lysis step, next to 200 μl homogenization buffer and 200 μl lysis buffer, 10 μl of a 20 mg/ml stock solution of proteinase K was added to each sample. Subsequently, samples were incubated for 10 minutes at 65 °C and 1000 rpm in a Thermomixer (Eppendorf, Hamburg, Germany). After cooling on ice for 1 minute, the standard protocol was followed.

### Sequencing

We used standard Illumina protocols for preparation and subsequent sequencing of RNA libraries. Libraries were sequenced on an Illumina HiSeq^™^ 2000 sequencing machine, using paired ends and 100 nucleotide read lengths. The raw data is available in the Sequence Read Archive (SRA; https://www.ncbi.nlm.nih.gov/sra) with ID PRJNA495983.

### SNP calling

The paired end reads were mapped against the N2 reference genome (WS220) using Tophat 49, allowing for 4 read mismatches, and a read edit distance of 4. SNPs were called using samtools 50 mpileup with bcftools and vcfutils.

### Construction of the genetic map

To construct a genetic map from the SNPs detected in the RNA-seq data we adjusted the method used in Serin & Snoek *et al.* 2017 51. For this *C. elegans* population we selected the SNPs by several parameters. First, we selected those SNPs present in at least one of the parental lines in both sequence replicates. Further selection was made based on i) presence in the mpRILs (min = 10, max = 180), quality (> 199), ii) correlation with neighboring SNPs of the same parental origin (> 0.8) and iii) heterozygosity (< 40 mpRILs). These SNPs (Supplemental Table 2) were used directly in the SNP map of the population (Supplemental Table 3) or translated to the parental origin genetic map (Supplemental Table 4).

## Phenotyping

### Population growth

Orsay/Santeuil mpRIL population growth was measured as the total offspring of 3 L4 hermaphrodites after 5 days at 20 °C in liquid peptone free medium (PFM). 24-well plates were inoculated with 1 ml liquid PFM per well and food bacteria added to a final OD_600_ of 5. The six different bacterial treatments were (i) *Escherichia coli OP50*, (ii) *Erwinia rhapontici* (isolated from Orsay, France), (iii) *Sphingobacterium sp.* (isolated from Orsay, France), (iv) a non-pathogenic *Bacillus thuringiensis* strain DSM-350E, and a pathogenic *Bacillus thuringiensis* strain NRRL B-18247 in the two concentrations of (v) 1:300 and (vi) 1:600. After 5 days, worms were fixed in 4 % formaldehyde and stored at 8 °C until counting 8.

### Lifespan assays

Worm lifespan assays were performed at 20 °C, with populations initiated from synchronized larvae isolated by incubating eggs from sodium hypochlorite treated gravid adults on plates without a food source 52. After 24 hours, the plates were seeded with *Escherichia coli* OP50 as a food source and the worms were allowed to grow, *en masse*, for 48 hours to the L4/young adult’s stage. After 48 hours, *ad libitum* (normal lifespan) worms were moved to fresh seeded standard NGM plates (5 worms per plate and 8 plates per treatment). The lifespan under DR worms were moved to seeded PFM plates (5 worms per plate and 8 plates per treatment). The method of total withdrawal of peptone from the agarose plates is a relatively mild form of DR, as described by Stastna *et al*. 201541. To test lifespan, worms were observed daily, with nematodes transferred to new plates every day until reproduction had ceased as assays were performed without the use of FUdR. After the reproductive period, the DR worms were moved to fresh plates every other day to prevent food deprivation. Worms were considered to have died if they were not moving and failed to respond to touch with a worm pick. Any worms that died due to maternal hatching (bagging) were censored out of the analysis of lifespan. Each mpRIL within an experimental block was tested at the same time under both conditions, with a total of 40 worms per treatment per mpRIL, plates were then randomized and blind coded. The movement of the L4/young adult worms to fresh plates was counted as day one for all the lifespan measurements. In total, the mpRILs were assayed in six blocks with 35-48 formally randomly selected mpRILs in each block, with some mpRILs present in multiple blocks. RILs were not included in the analysis if the lifespan of less than three worms was observed per treatment. In addition to the mpRILs, the four parental lines and N2 were also tested in all lifespan assays.

### Heat shock resistance

Worms were cultured at 15 °C prior to the heat shock assays. Worms were synchronized as for the lifespan assay and allowed to grow *en masse* to L4/young adult stage [52-54]. At this stage, worms were transferred to fresh plates, 10 worms per plate with five replicates for each of the mpRILs and each of the parental strains. The plates were then randomized and blind coded. Worms were then placed at 35 °C for 10 hours. After the heat shock, worms were allowed to rest at 15 °C for 48 hours before scoring for survival, when worms that did not respond to a gentle prod with the worm pick were scored as dead. Worms that crawled off the plates or died of bagging were censored from the experiment. The data were then converted into a proportion of survival.

### Oxidative stress resistance

Worms were maintained at 20 °C, synchronized as described above and grown *en masse* to the L4/young adult’s stage. After 48 hours, the worms were washed off the plates with M9 buffer and 10-30 individuals were transferred to 96 well plates in a total volume of 48 μl, with three replicates for all the mpRILs and N2. The plates were then transferred to a WMicrotracker-One™ (PhylumTech) and activity over 30 minutes was determined at 20 °C. After this step, 2 μl of 0.4% H_2_O_2_ solution was added to all wells, giving a final volume of 50 μl, except for the control, which had 2 μl of M9 buffer added to make up the final volume. Worms were then incubated for 24 hours at 20 °C. After 24 hours, the locomotive activity of the worms was measured again. WMicrotracker-One^™^ records movement as photo-beam interruptions within wells of 96-well plates. The data were then processed as follows, (activity of the wells before - activity after 24 hours)/before = activity score. An activity score of -1 represents no movement and hence that all worms were dead at the end of the treatment, and an activity score of 0 indicates an activity level after 24 hours that is the same as before. This score can also generate values above zero, which indicates that worms were more active after the hydrogen peroxide treatment.

### Developmental time and occurrence of males

Starvation synchronized L1 juveniles were grown on *E. coli* OP50 at 24 °C and after 48 h inspected at 1 hour time intervals. Developmental time was defined as the period between synchronized hatching and time till first eggs. Time till first egg scoring was adjusted from [8, 55] by placing 20-40 worms on NGM, done *in duplo*, and scoring every hour starting at 48 h till 54 h for eggs. This was done on *E. coli* OP50 and *Erwinia rhapontici*, previously 8. We scored time of first egg visible on plate and time when multiple groups of ~10 eggs were visible. Averages of these time points per mpRIL were used in QTL mapping. Moreover, the occurrence of males on the plates was recorded after population growth and used for QTL mapping.

### Size and volume

Analysis of length and width of young gravid adults was performed with a particle analyzer (RapidVue; Beckman Coulter Inc., Miami, FL, USA)8.

## QTL mapping

We started with single marker mapping for each trait to find the SNP with most significant QTL (Supplemental Figure 4). This SNP was used as the starting point in the forward mapping approach. Forward mapping was done by selecting cofactors one by one, starting with the most significant and remap with that cofactor and selecting the next most significant SNP till no more SNP was present with a p < 0.001 or a maximum of 10 cofactors was reached. Then QTLs were remapped with the selected cofactors and an exclusion window of 2 Mbp. Cofactors within this window were excluded from the mapping model when QTLs were mapped in the window. Obtained QTLs were determined significant when-log10(p) > 3 and borders were determined at the point where the QTL profile drops 2-log10(p) scores below the peak. (permutations showed maximum QTLs ranging from-log10(p) of 2.1 to 3.9, with the exception of the traits describing the occurrence of males on the plate for which in a number of permutations a-log10(p) was found > 4.5). All QTL profiles are can be obtained and interactively explored in EleQTL (http://www.bioinformatics.nl/EleQTL). Heritability for each trait was calculated by dividing the variation between the mpRILs by the total variation.

## Supplemental files

**Supplemental Figure 1: Distribution of parental alleles in the multi-parental Recombinant Inbred Lines**. Colours indicate the percentage parental occurrence per genomic position (x-axis) per cross (y-axis) as estimated from the parental SNP Distribution Patterns (SDP). Chromosomes are in separate panels on the x-axis. mpRILs are grouped according to their cross history. Group Z are the parental lines.

**Supplemental Figure 2: SNP Distribution Pattern (SDP) per mpRIL.** For each of the 7 SDPs the genotype for each mpRIL is shown. SDPs are shown on top, chromosomes on the right. The genotype of the mpRIL is green when it is has the SNP corresponding to the SDP and red when it has the opposite variant. SDP 12: (JU1511/JU1926 vs JU1931/JU1941), 13: (JU1511/JU1931 vs JU1926/JU1941), 14: (JU1511/JU1941 vs JU1926/JU1931), JU1511: (JU1511 vs rest), JU1926: (JU1926 vs rest), JU1931: (JU1931 vs rest) and JU1941: (JU1941 vs rest). Notice that each recombination event can break up multiple SDP.

**Supplemental Figure 3: Forward mapping QTL profiles for each trait.** Trait names shown on the right. Chromosome number shown on top. Genomic position in Mbp shown on the x-axis. For each SNP the significance in – log10(p) multiplied by the sign of the effect on the y-axis. Colours indicate SPD of the SNP and triangle the cofactors used in the final model of the forward mapping approach.

**Supplemental Figure 4: Single marker QTL profiles for each trait.** Trait names shown as title. Chromosome number shown on top. Genomic position in Mbp shown on the x-axis. For each SNP the significance in – log10(p) multiplied by the sign of the effect on the y-axis. Colours indicate SPD of the SNP.

**Supplemental Table 1: Detailed crossing scheme used to make the mpRILs.** The crosses from which each individual mpRIL was made can be found here.

**Supplemental Table 2: SNP info.** SNP position and SNP distribution pattern (SDP)

**Supplemental Table 3: SNP genetic map.** SNP identity per mpRIL

**Supplemental Table 4: Parental background genetic map.**

**Supplemental Table 5: Average phenotypic values per line used for QTL mapping.**

**Supplemental Table 6: Trait descriptive.** Number of mpRILs for which the phenotype was measured, minimum trait value, maximum trait value, mean trait value, median trait value, trait value of parental line JU1511, trait value of parental line JU1926, trait value of parental line JU1931, trait value of parental line JU1941, heritability, heritability type, number of QTLs found and explained variation by QTLs.

**Supplemental Table 7: Correlation between Traits.**

**Supplemental Table 8: Identified QTLs.**

